# Beyond Immobility: Computational Modeling Reveals Cognitive Processes in Simple Rodent Depression Tests

**DOI:** 10.1101/2025.01.24.634822

**Authors:** Zhihan Li, Tianyu Lu, Jiaozhao Yan, Xiang Zhang, Yun-Feng Li

**Affiliations:** Beijing Institute of Basic Medical Sciences, Haidian, Beijing,100039, China; School of Life Science and Technology, ShanghaiTech University, Pudong, Shanghai, 201210, China; School of Pharmaceutical Sciences, Anhui Medical University, Hefei, Anhui, 230032, China; Beijing Institute of Pharmacology and Toxicology, Haidian, Beijing, 100039, China

**Keywords:** forced swim tests, tail suspension tests, depression-like behaviors, cognitive processes, computational modeling, reinforcement learning models

## Abstract

Simple behavioral tests like the forced swim test (FST) and tail suspension test (TST) are widely used to assess depression-like behaviors in rodents, primarily measuring immobility time. However, this approach oversimplifies behavioral readouts and overlooks the cognitive processes driving behavior, leaving the relationship between increased immobility and cognitive biases unclear. Here, we developed the SwimStruggleTracker (SST) to extract fine-grained behavioral trajectories and integrate computational modeling to methodically analyze behavior. Our findings reveal that behavior in the FST and TST follows reinforcement learning principles involving learning, consequence perception, and decision-making. Notably, the cognitive processes underlying behavior differ between the two tests, challenging the assumption that they are interchangeable for cross-validation. Regression analyses identify distinct behavior phases: early behavior is primarily influenced by learning-related factors, while later stages are more affected by consequence sensitivity. These findings suggest traditional analyses focusing final minutes may underestimate the role of learning and overemphasize consequence sensitivity.

**Motivation:** The forced swim test (FST) and tail suspension test (TST) are among the most widely used paradigms for assessing depression-like behaviors in rodents. Yet, traditional analyses typically quantify only immobility during the final minutes, discarding rich temporal structure in the data and hindering efforts to uncover the cognitive mechanisms underlying these behaviors. To address this gap, we developed an automated tool that captures behavioral trajectories with fine temporal resolution and integrates computational modeling to dissect the cognitive processes driving behavior. Using this approach, we demonstrate that the FST and TST engage overlapping but partially distinct cognitive processes, and that the dominant cognitive components shift across different stages of the tests.

**Highlights:**

- SwimStruggleTracker (SST) accurately rejects passive movements, such as pendulum-like motion.
- Reinforcement learning models capture the behavioral dynamics of mice in the FST and TST.
- Distinct winning models indicate that the FST and TST engage partially dissociable cognitive processes.
- Learning factors dominate early stages, whereas consequence-sensitivity factors dominate later stages.

## Introduction

Recent advances in rodent research have substantially expanded our understanding of the neural circuits and molecular mechanisms underlying mental disorders, establishing this field as a cornerstone of neuroscience and psychiatry^1–5^. However, rodents’ inability to verbally express subjective experiences or perform complex cognitive tasks limits their ability to capture the cognitive distortions and information-processing abnormalities seen in conditions like depression^6–8^. To overcome this limitation, simplified behavioral models—including the forced swim test (FST) and tail suspension test (TST)—have been widely adopted^9–13^. Over the past two decades, these methods have seen a marked increase in usage and remain among the most popular behavioral assays (Figure 1A-D). The primary measure in both tests is immobility time, often interpreted as an indicator of depression-like behavior. Yet, this narrow focus on immobility overlooks the underlying cognitive mechanisms, raising questions about what these tests truly measure. One promising explanation, the behavioral adaptation hypothesis, proposes that the gradual increase in immobility reflects an adaptive learning process^9,14–19^. Despite its prominence, empirical evidence supporting this hypothesis remains sparse, underscoring the need for further investigation into the cognitive processes driving behavior in these tests.

**Figure 1.**
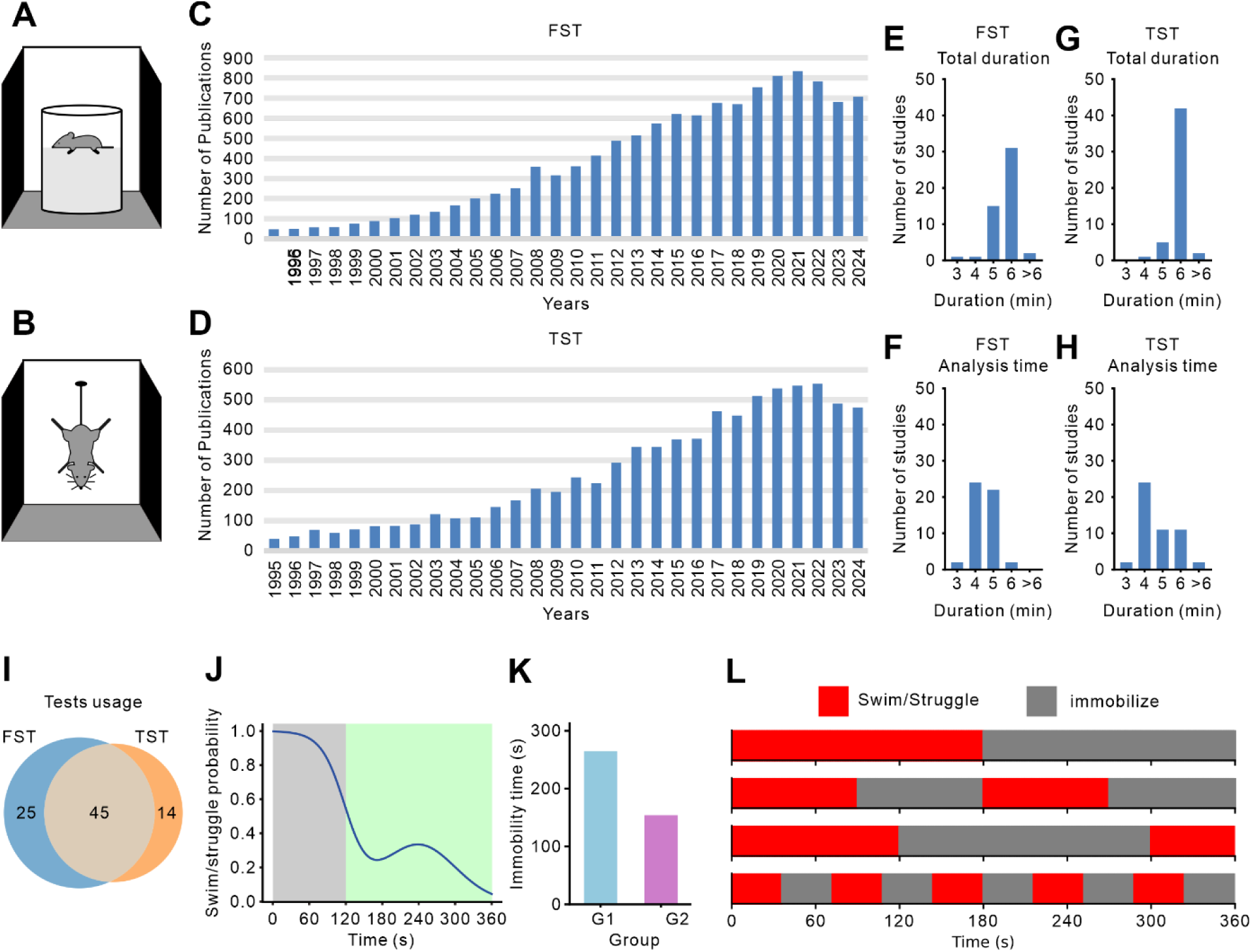
The forced swim test (FST) and tail suspension test (TST). (A, B) Experimental setup: Both tests subject mice to inescapable conditions for approximately 5-6 minutes. In the FST (A), mice are placed in a water-filled container and behavior is categorized as either swimming or immobility. In the TST (B), mice are suspended upside down and behavior is classified as struggling or immobility. (C, D) Research trends. Data from Web of Science over the past two decades shows a steady increase in publications employing the FST (C) and TST (D) which underscores their growing prominence in behavioral research. (E-H) Test duration and analysis time window. We reviewed 50 recent studies using the FST (E, F) and 50 using the TST (G, H), recorded both the total test duration and the specific analysis window. Most studies focused on the final 4-5 minutes rather than the entire test period. (I) Joint application of FST and TST. After excluding 16 duplicate studies, we assessed the concurrent use of both tests, finding that the majority of studies employed them together in their experimental design. (J) Averaged behavioral trajectory. This schematic depicts a typical decline in swimming or struggling behavior over time with fluctuations in the later stages. Traditional analyses often emphasize the final minutes (green background) while neglecting the early phases (grey background). The data in this panel are artificially generated for demonstration. (K) Conventional behavioral analysis. Traditional methods typically use total immobility time as a metric. For illustrative purposes only, artificial data are used, with G1 and G2 representing two hypothetical experimental groups. (L) Individual behavioral trajectories. The cartoons highlight the complexity of behavioral patterns and demonstrate that even if a mouse remains immobile for half of the 6-minute test, distinct behavioral trajectories may emerge.

A key challenge in elucidating these cognitive processes lies in the limited scope of current behavioral metrics. Most studies condense complex behaviors, typically recorded over 5-6 minutes, into a single measure of total immobility time in final minutes (Figure 1E-K). This simplification obscures potentially informative behavioral patterns that could illuminate underlying cognitive mechanisms. For instance, as illustrated in Figure 1L, animals with the same overall immobility times may exhibit markedly different behavioral trajectories, suggesting engagement of distinct cognitive processes. Although some pioneering studies have examined high-resolution behavioral trajectoires^10^, they often rely on labor-intensive manual annotations, limiting scalability and broader application. Consequently, there is a growing demand for automated tools capable of efficiently capturing and analyzing fine-grained behavioral data, thereby advancing research in this area.

Although open-source frameworks such as DeepLabCut (DLC)^20–23^ and LEAP^24,25^ have significantly advanced animal pose estimation, they typically output the positions of animal’s keypoints, leaving researchers to develop separate classification models to translate these poses into behaviors like immobility, swimming, or struggling. Moreover, they ofen require manual annotation of training data for fine-tuning before any inference can be performed, which complicates application. Commercial packages (e.g., SMART, EthoVision) offer end-to-end behavioral detection but are prohibitively costly for many laboratories. As a result, whether due to financial constraints or underappreciation of the rich information encoded in detailed trajectory data, researchers frequently forego these solutions. Collectively, these barriers severely restrict the automated extraction of fine-grained behavioral metrics in FST and TST.

In addition to improving data collection, a further challenge is interpreting these data to uncover the cognitive mechanisms at play. Computational modeling offers a powerful framework for formalizing latent cognitive computations that underlie behavioral patterns, with growing applications in psychiatric research^26–31^. By enabling quantitative decomposition of behavior sequences across various experimental paradigms, such as probabilistic reversal learning (PRL)^32–35^, Iowa gambling task (IGT)^36^, fear conditioning^37^ and two-stage decision^38,39^, this approach has revealed cognitive differences among distinct subject groups, particularly regarding learning dynamics^40,41^, prediction errors^42–44^, and perception of rewards and punishments^45–50^. Such insights have refined cognitive models of mental disorders, offering a basis for improved understanding and potential therapeutic development. Although computational methods like reinforcement learning models are increasingly applied to rodent behavior^51–55^, current studies have focused mainly on operant learning tasks. Consequently, despite their widespread use, the FST and TST remain underexplored with respect to the cognitive mechanisms and information-processing biases that underlie depressive-like behaviors, which we aim to characterize in this study.

In this study, we developed a custom tool to extract fine-grained behavioral trajectories from the FST and TST in mice and applied reinforcement learning models to infer the underlying cognitive processes. Our analysis shows that behavior in the FST and TST is best captured by distinct reinforcement learning models, suggesting that these assays engage partially dissociable cognitive processes. Furthermore, regression analyses of model parameters reveal that different stages of the tests are governed by distinct cognitive factors. These findings highlight a potential limitation of conventional analysis approaches, which often overlook early behavioral responses and may therefore underestimate learning-related processes while overemphasizing consequence sensitivity.

## Results

### An open-source tool for annotating behavioral trajectories

Detailed behavioral trajectories are crucial for analyzing the cognitive processes underlying behavior. However, manual annotation is time-consuming and subject to individual biases, potentially compromising both accuracy and consistency. To address these challenges, we developed an automated tool, SwimStruggleTracker (SST), designed to annotate fine-grained behavioral states in the FST and TST^1^. As shown in Figure 2, the process begins with calibration and region of interest (ROI) selection, which corrects for image distortion due to camera angles and allows users to define the region where mouse movement is expected, termed the ROI. Next, the passive motion elimination module filters out irrelevant movements, such as floating or swaying, which could otherwise result in false detections. The active motion detection module then identifies the mouse’s state, whether it is swimming, struggling, or immobile, on a frame-by-frame basis. Finally, the behavioral trajectory smoothing and binning module smooths the data and bins it into 2-second intervals, producing a clean and reliable behavioral trajectory for further analysis and modeling.

**Figure 2.**
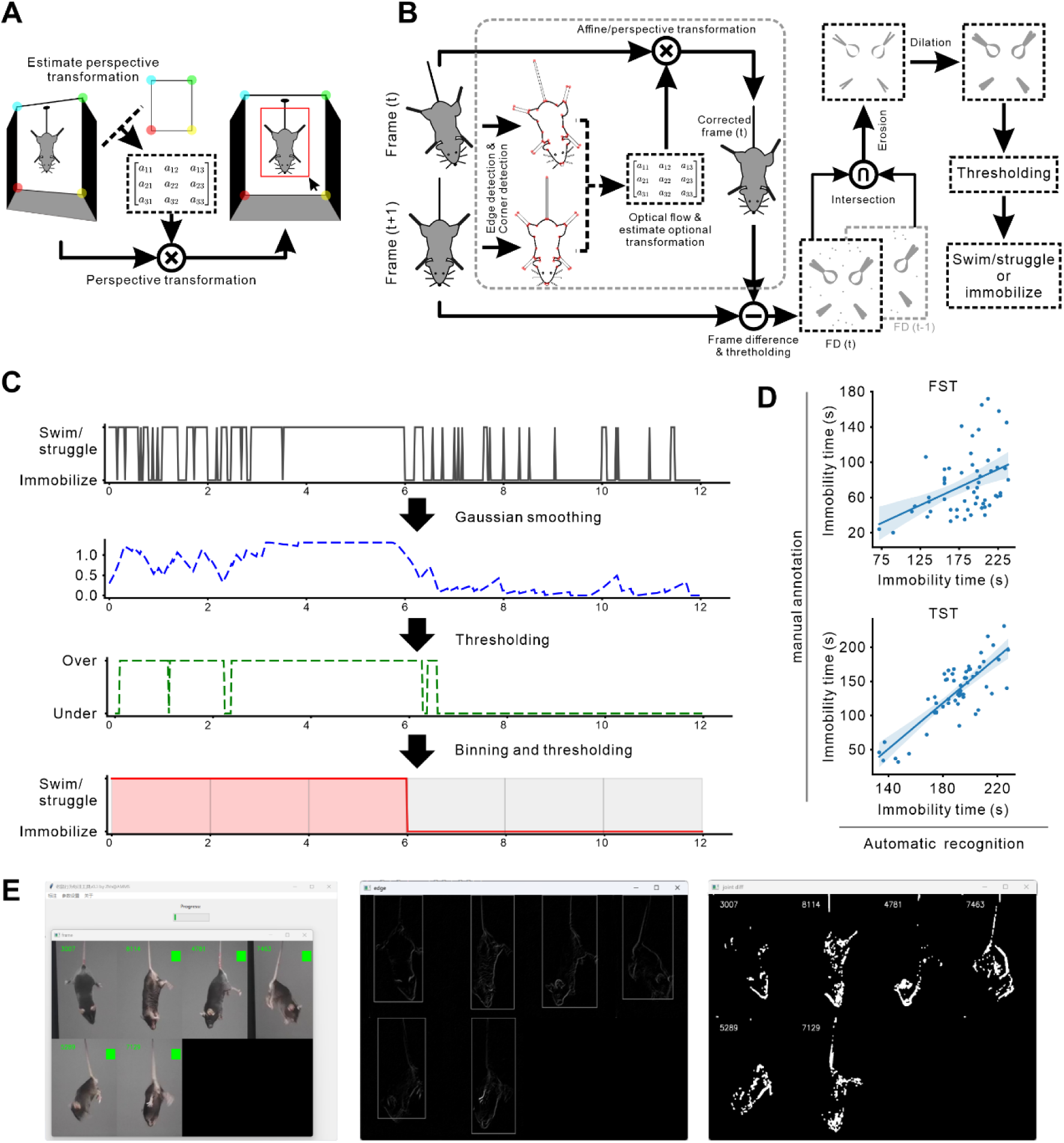
Overview of the SwimStruggleTracker (SST). (A) Frame calibration and ROI selection. Raw video frames are corrected through perspective transformation based on camera angles. The user manually selects four vertices of the experimental box, which are compared to pre-measured dimensions to generates a transformation matrix that corrects the frames. The region of interest (ROI) is defined as the maximal area available for mouse movement. (B) passive motion elimination and active motion detection. The SST identifies behavioral states by calculating differences between consecutive frames. An optional module removes passive motion (e.g., floating or swinging) using geometric transformations to enhance accuracy. Noise is reduced through morphological operations, followed by refinement of the trajectory using smoothing, binning, and thresholding techniques. (C) Smoothing and binning of behavioral trajectories. Raw behavioral trajectories can be noisy, such as when a 6-second struggle is followed by 6-seconds of immobility, resulting in erratic data. A Gaussian smoothing step assigns a continuous behavioral value to each frame. Thresholding then converts the behavioral values into binary profiles, which are subsequently binned into 2-second intervals to produce a final, clear trajectory. (D) Validation with human comparison. We validated SST’s performance by comparing its outputs with manual annotations. (E) User interface. The user-friendly interface enables efficient operation and real-time visualization of intermediate outputs, facilitating precise monitoring of each key processing step.

We validated SST’s performance by comparing its outputs with human expert annotations. Significant correlations observed in both total immobility times (Figure 2D, FST: r = 0.411, p = 0.002; TST: r = 0.823, p < 10^−5^, Pearson correlation), indicating close agreement between SST and manual scoring. To further evaluate SST, we asked human experts to adjudicate discrepancies between SST and the widely used commercial software SMART v3.0 on TST videos from five mice. Among segments with conflicting annotations, 77.8% of the segments (63 vs. 18) and 89.5% of the total disputed time (436s vs. 56s) were judged in favor of SST (Figure S1). Notably, SST demonstrated strong robustness in correctly rejecting pendulum-like passive movements, a common artifact in the TST that is frequently misclassified as active behavior by other systems (see Supplementary Video). All subsequent analyses will be based on data provided by SST.

### Behavioral trajectories in the FST and TST

To induce depression-like behaviors, we employed two experimental manipulations: chronic restraint stress^56^ (CRS; Figure 3A) and chemogenetic inhibition of medial prefrontal cortex (mPFC) pyramidal neurons^57–61^ (Figure 3H). Average behavioral trajectories for CRS are presented in Figure 3B and C, while those for mPFC inhibition are shown in Figure 3I and J. Notably, the probabilities of swimming and struggling typically declined after test onset, with swimming showing a rebound primarily in the FST, whereas late-stage variability increased in both the FST and TST. In the FST, both CRS and mPFC inhibition lead to reduced swimming behavior in the later stages compared to controls (Figure 3B, I). In the TST, CRS produced a reduction in struggling behavior during the early stages (Figure 3C). Conventional analyses of immobility times corroborate these findings and align with the existing literature (Figure 3D-G, K-N), supporting the reliability of our behavioral trajectory data. Furthermore, the CRS group exhibited significantly shorter inter-struggle-intervals in both the FST (Figure S2E) and TST (Figure S2G). In the TST, CRS also reduced struggle bout duration (Figure S2K), with a similar trend observed in the FST (Figure S2I). By contrast, the hM4D group showed only a marginal decrease in inter-struggle interval relative to controls (Figure S2F).

**Figure 3.**
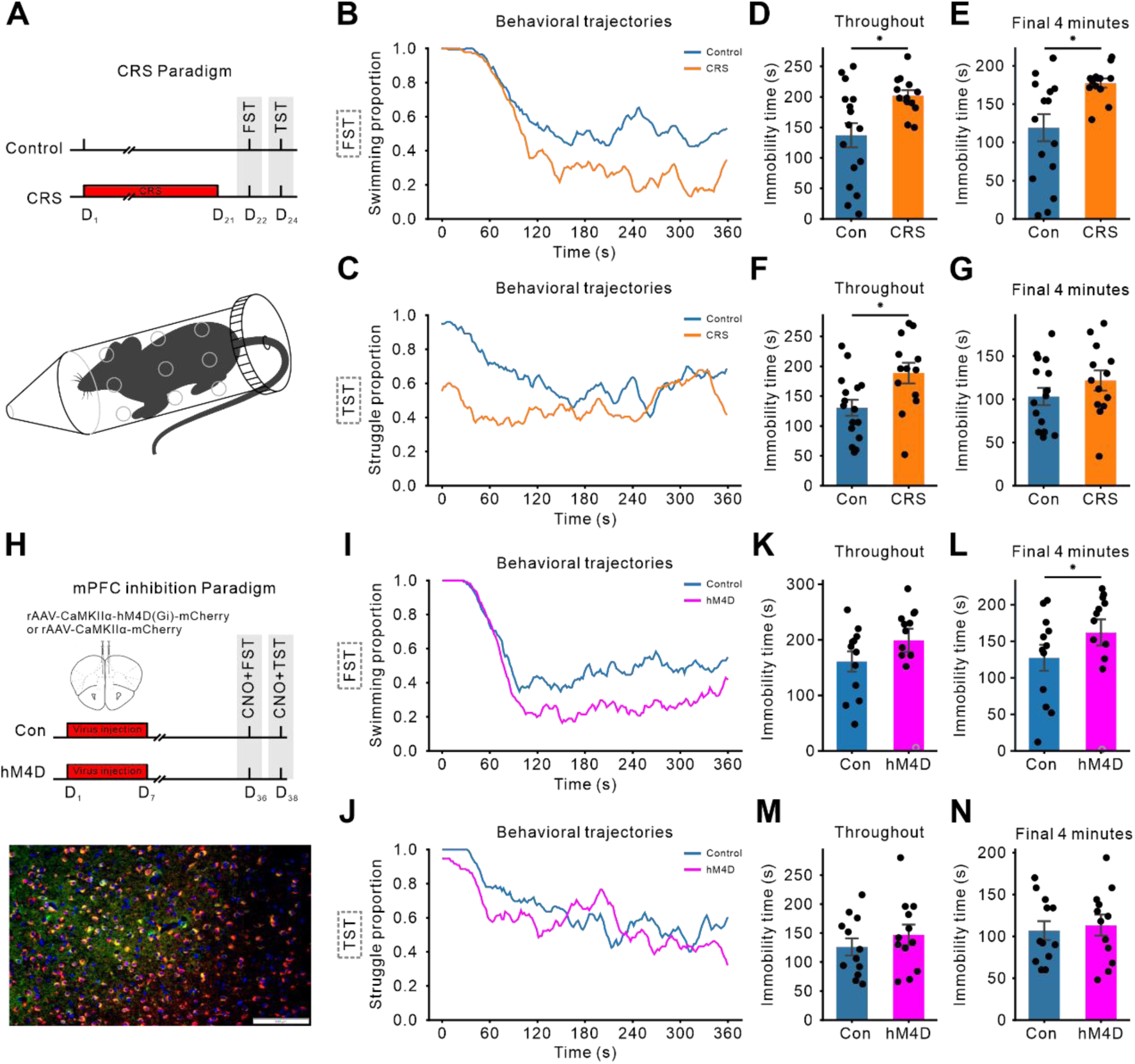
Behavior analysis of mice in FST and TST under two experimental manipulations. (A) Experimental setup for CRS. Twenty-nine C57BL/6 mice were divided into two groups; the CRS group (n = 13) underwent daily restraint stress (6-8 hours/day) for 21 days, while controls (n = 16) experiencing no stress. Behavioral assessments were conducted on days 22 (FST) and 24 (TST). (B, C) Behavioral trajectories of CRS and control mice during the FST and TST. Averaged trajectories highlight stress-induced behavioral changes, with control mice shown in blue and CRS mice in orange. (D-G) Immobility times for the FST (D, E) and TST (F, G) in the CRS experiment are shown for the entire testing period (D, F) and the final 4 minutes (E, G). Blue and orange bars represent the control and CRS group, respectively, while black dots indicate individual animal data. Error bars denote the standard error of the mean (SEM). (H) Chemogenetic inhibition of mPFC pyramidal neuron. Immunofluorescence confirmed hM4D(Gi)-mCherry expression in mPFC pyramidal neurons (red=hM4D(Gi), green=CaMKIIα, blue=DAPI; scale bar = 100 μm). Viral expression colocalized with pyramidal neurons in 89.27% of cells (n=3 mice). (I, J) Behavioral trajectories for mPFC inhibition in the FST and TST, with blue curves representing controls and magenta curves indicating hM4D-treated mice, demonstrating the effect of mPFC inhibition. (K-N) Immobility times for the FST (G, H) and TST (I, L) in the mPFC inhibition experiment follow the same format as panels (D-G). Blue and magenta bars represent the control and hM4D groups, respectively. Hollow gray circles in panels (L, N) denote the outlier mice that swam for nearly the entire test. Asterisks denote significant differences from the controls as determined by the Mann–Whitney U test; **p < 0.01, *p < 0.05.

These detailed behavioral trajectories provide a richer dataset for nuanced analysis, as their dynamic patterns likely reflect underlying cognitive processes in mice, Building upon this data, we employ computational modeling to elucidate the cognitive processes driving depression-like behavior.

### Behaviors in the FST and TST are driven by different reinforcement learning processes

The observed behavioral trajectories exhibit patterns consistent with adaptive behavior, motivating the use of reinforcement learning as a computational framework to model these dynamics (Figure 4A). We evaluated eight candidate models (see Methods for details), including the classical Rescorla-Wagner (RW) and Pearce-Hall (PH) models, along with their extensions incorporating three additional cognitive components: perseveration (+per), consequence sensitivity (+rho), and adaptation (+ada). Model comparison was based on model weights, representing the relative likelihood that a given model was the true generative process under each experimental condition. The model with the highest weight was identified as the winning model, providing the best account of the observed data.

**Figure 4.**
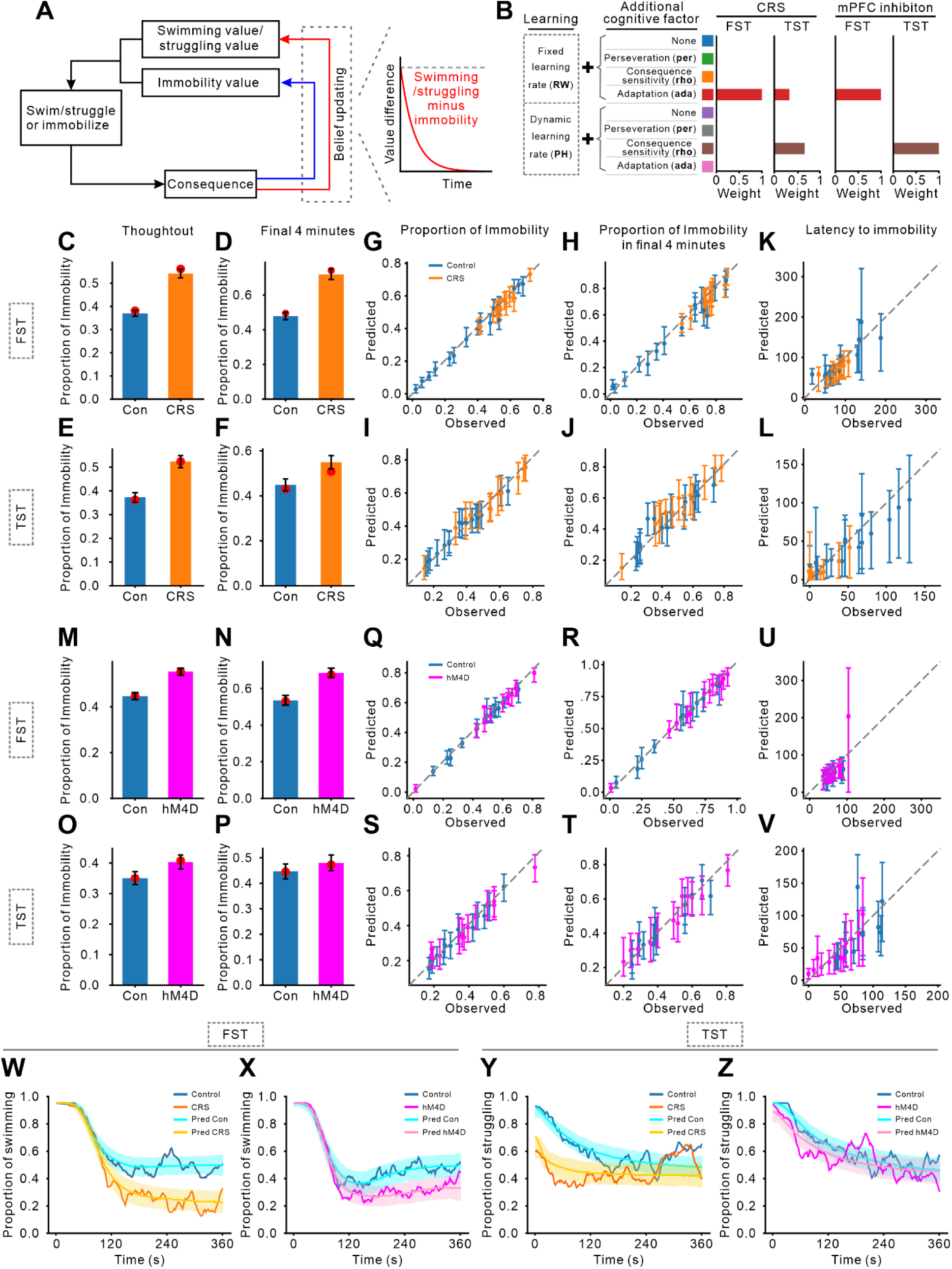
Comparing and validating reinforcement learning models. (A) Reinforcement learning framework. Mice continually evaluate expected consequence of two actions—swimming vs. immobility in the FST and struggling vs. immobility in the TST—and update these expectations based on environmental feedback. Decision-making is guided by comparing the expected consequence of each action. (B) Model Comparison. The left side of the panel illustrates the learning rules and cognitive components included in each model. The bar plots display model weights for each test and experimental condition, with different colors representing distinct models. Across conditions, the winning models were RW+ada in the FST and PH+rho in the TST. (C-F) Model-predicted vs. observed proportions of immobility (CRS experiment). Panels (C, D) display predictions for the FST, and (E, F) for the TST. Each pair compares model predictions with observed data over the full testing period (C, E) and the final 4 minutes (D, F). Colored bars indicate median predictions from the winning models, with error bars denoting the 95% highest density interval (HDI). Red dots present the observed group means, showing close alignment between model predictions and empirical data. (G-J) Individual-level predicted vs. observed proportions of immobility (CRS experiment). Panels (G, H) show results for the FST, and (I, J) for the TST. The x-axis represents observed values, and the y-axis indicates median model predictions based on 2000 simulations. Error bars denote the 95% HDI, and different colors denote experimental groups. (K, L) Predicted vs. observed latencies to immobility (CRS experiment), using the same format as in panels (G–J). (M-P) Model-predicted vs. observed proportions of immobility (mPFC inhibition experiment), replicate the analysis from (C-F) for the mPFC inhibition experiment. (Q-V) Individual-level predicted vs. observed immobility proportions and latencies (mPFC inhibition experiment). These panels replicate the individual-level analyses from (G–L) for the mPFC inhibition experiment. (W-Z) Predicted versus observed behavioral trajectories. Panels (W, X) show model predictions for swimming behavior in the FST, while panels (Y, Z) display predictions for struggling behavior in the TST. Predicted trajectories are presented by cyan, yellow, and pink curves (median predictions), with shaded areas representing 95% highest density intervals (HDI) from 2000 simulations. Observed data are overlaid as blue, orange, and magenta lines, demonstrating strong concordance between model predictions and experimental observations.

As shown in Figure 4B, the winning models differed between the FST and TST. In the FSTs, the winning models were RW+ada, which updated beliefs with a fixed learning rate while dynamically adjusting sensitivity to the consequences of actions. In contrast, the winning models in the TSTs were PH+rho, which modulated the learning rate based on attention allocation—shaped by discrepancies between perceived and expected consequences—while maintaining a fixed sensitivity to outcomes. Both models successfully recovered their parameters (RW+ada: r>0.52, p<0.001; PH+rho: r>0.64, p<10^−4^; Pearson correlation), confirming their validity and reliability in capturing the behavioral data.

To evaluate the predictive power of these winning models, we conducted posterior predictive checks. As shown in Figure 4C-F and M-P, both models successfully captured the observed immobility proportions across all groups, both over the entire task and during the final 4 minutes. Nearly all individual immobility times fell within the predicted 95% highest density interval (HDI, Figure 4G-J and Q-T), with strong correlations between predicted and observed immobility times across all experimental conditions and time windows (r>0.95, p<10^−5^, Pearson correlation). Figure 4K, L, U, and V demonstrate that the models reliably predicted each mouse’s latency to immobility (r>0.71, p<0.001, Pearson correlation). Moreover, the winning models accurately replicated the mice‘s behavioral trajectories (Figure 4W-Z), capturing the dynamic evolution of behavior over time. Collectively, these results confirm that the winning models robustly simulate mouse behavior.

These findings indicate that the evolution of behavior in both the FST and TST can be effectively explained by reinforcement learning principles, reflecting ongoing processes of learning, consequence perception, and decision-making. Notably, the distinct winning models for these tests suggest that different cognitive processes drive behaviors in the FST and TST. This challenges the prevailing assumption that the FST and TST are interchangeable for cross-validation, highlighting the need to account for task-specific cognitive processes.

### CRS and mPFC inhibition increase immobility by altering learning and consequence sensitivity

Within the framework of reinforcement learning, model parameters represent distinct cognitive components, thereby clarifying how specific factors contribute to increased immobility. In the FST (Figure 5A, B), both the CRS and hM4D groups demonstrated significant reductions in the learning rate (α) and the lower bound of consequence sensitivity adaptation (lb), alongside an elevated adaptation rate (β) compared to the controls. Additionally, The hM4D group exhibited a higher inverse temperature (τ). In the TST (Figure 5C, D), both CRS and hM4D groups displayed lower τ values than controls, while the CRS group further demonstrated increased scaling factor (κ) and attention updating rate (η), together with decreased consequence sensitivity (ρ).

**Figure 5.**
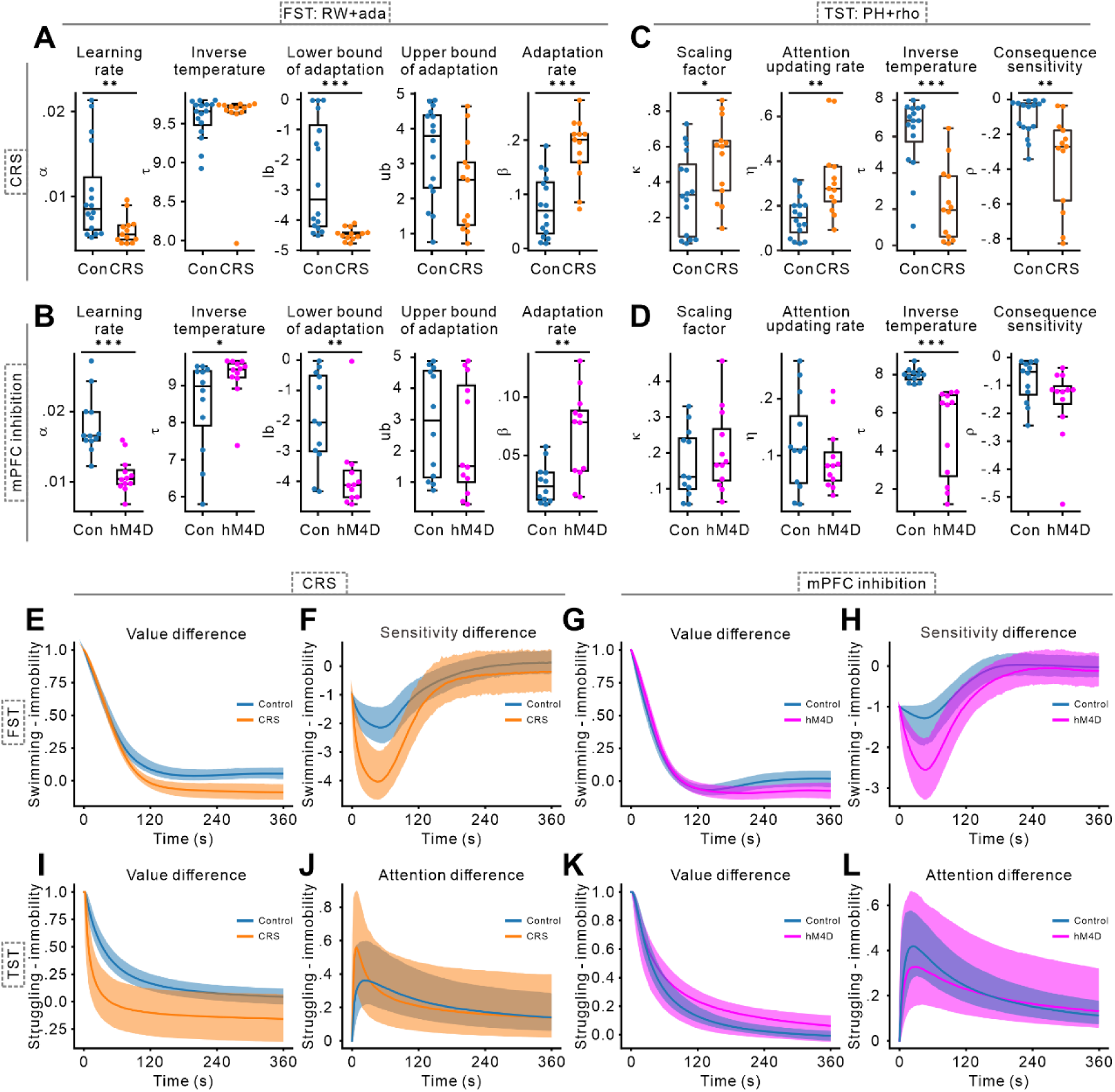
Uncovering cognitive processes driving immobility via model parameters. (A, B) Parameter distributions of the RW+ada model for the FST. This model comprises five parameters: the learning rate (α), which determines how rapidly animals update expectations; the inverse temperature (τ), which regulates the impact of expected values on decision-making; the lower bound of adaptation (lb), reflecting intensified negative experiences from repeated failed escape attempts; the upper bound of adaptation (ub), representing diminished negative perception after sustained immobility; and the adaptation rate (β), which dictates the speed of adaptation. Individual data points representing the mean of 2000 samples per mouse, with group-level medians and interquartile ranges depicted by boxplots. Control, CRS, and hM4D groups are represented in blue, orange, and magenta, respectively. (C, D) Parameter distributions of the PH+rho model for the TST. This model includes four parameters: the scaling factor (κ), which modulates the influence of attention on learning; the attention update rate (η), governing the pace of attentional learning; τ, which functions similarly to that in the RW+ada model by guiding decision-making; and consequence sensitivity (ρ), which quantifies the perceived severity of failure following unsuccessful struggles. Panels formatting is consistent with (A, B). (E-H) Predicted dynamics of expected value and consequence sensitivity in the FST. Panels (E, F) show differences between swimming and immobility under CRS conditions, while panels (G, H) display results for mPFC inhibition. Solid lines represent group-level medians, and shaded areas indicate 95% HDI derived from 2,000 posterior samples. (I-L) Predicted dynamics of expected value and attention allocation in the TST. Panels (I, J) present struggling versus immobility under CRS conditions, whereas panels (K, L) present predictions for mPFC inhibition. Formatting is consistent with (E–H). Asterisks indicate significant differences compared to controls (***p < 0.001, **p < 0.01, *p < 0.05; Mann–Whitney U test).

We next examined intermediate variables including the expected value, adapted consequence sensitivity, and attention allocation throughout the tests (Figure 5E-L). Over time, the relative expected value of active behaviors (swimming/struggling) vs. immobility decreased and eventually plateaued at a lower level. In the FST, both CRS and hM4D groups showed reduced differences in late stages compared to controls (Figure 5E, G), whereas in the TST, the CRS group exhibited a more rapid early decline, with the hM4D group remaining similar to controls (Figure 2I). For relative consequence sensitivity, both the CRS and hM4D groups displayed lower values than controls in the early stages of the FST (Figure 5F, H). Regarding attention allocation in the TST, the CRS group devoted slightly more attention to struggling behavior in the early stage compared to the control group (Figure 5J).

Taken together, the decrease in lb in the FST suggests that CRS and mPFC inhibition may heighten despair after failed escape attempts. The increase in β likely accelerated this sense of despair, as evidenced by the steeper early decline of the orange and magenta curves in Figure 5F and H. Consequently, the expected value for swimming declines (Figure 5E, G), leading to higher immobility. In the TST, the early increase in η for the CRS group indicates that these mice more quickly recognized discrepancies between expected and actual outcomes of struggling. (Figure 5J, where the orange line is above the blue line in the early stages). Coupled with a higher κ, this accelerated the learning of the negative consequences associated with struggling (Figure 5I, where the orange line declines more rapidly), ultimately leading to an earlier onset of immobility.

### Differential roles of learning rate and consequence sensitivity in early and later behavior

To investigate how model parameters shape behavior, we conducted simulations in which two parameters were varied simultaneously while all others were held at their median values, derived from the posterior distributions of the winning models (Figure 6A-H). In the FST, higher lower (lb) and upper (ub) bounds of consequence sensitivity adaptation were associated with prolonged swimming (Figure 6B, D). The relationship between α and β was more complex: very low α values led to extended swimming, whereas moderate β values, when combined with higher α, promoted swimming behavior (Figure 6A, C). In the TST, longer struggling durations were linked to higher ρ and lower κ, η and τ (Figure 6E-H).

**Figure 6.**
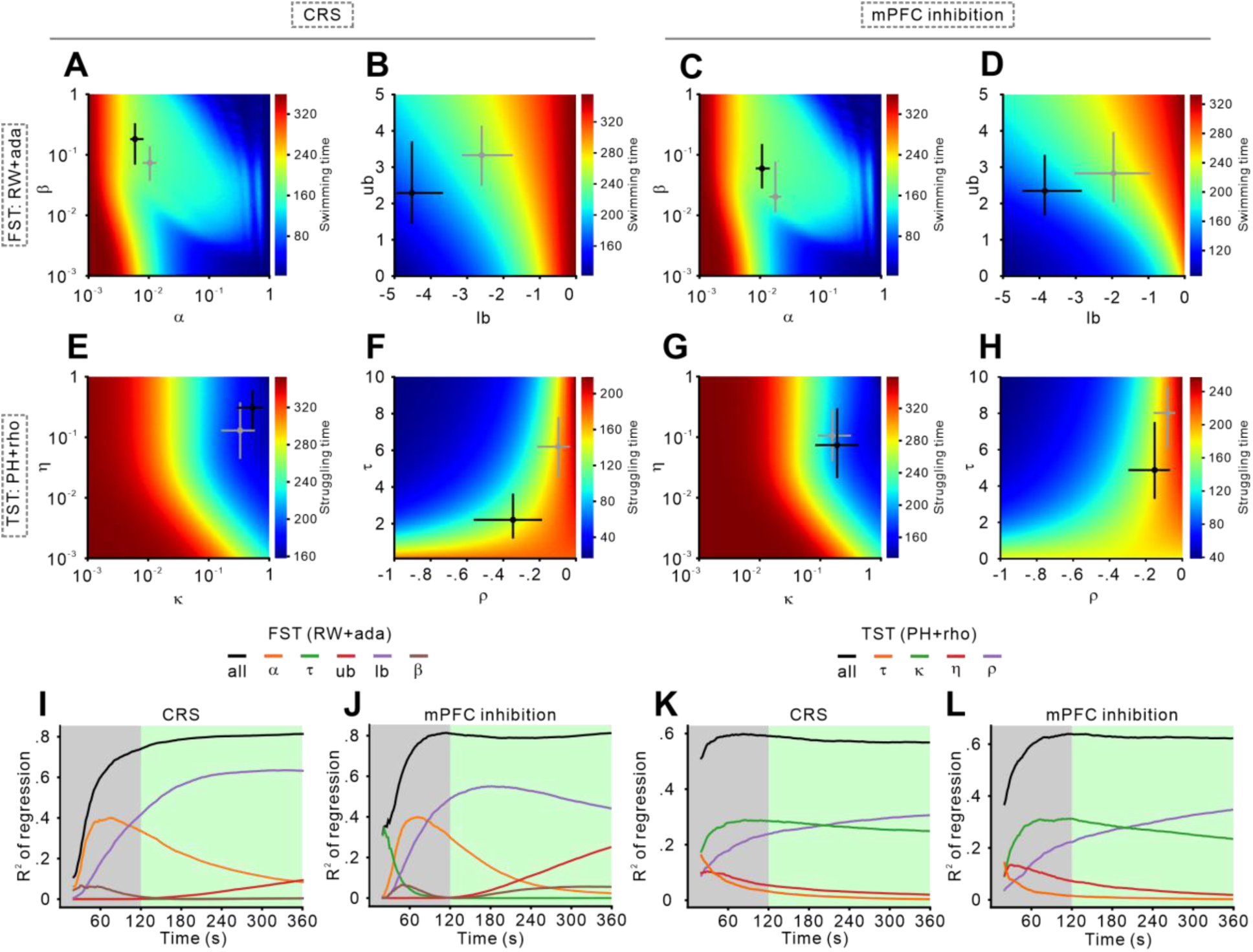
Linking model parameters to behavioral patterns. (A-D) Simulated swimming durations with the RW+ada model. Heatmaps illustrate total simulated swimming times across the parameter space of the RW+ada models. Each heatmap explores the effects of two parameters while others remain fixed at their posterior median. The x- and y-axes denote the sampled parameter values, with brighter colors indicating longer swimming durations. Circles at the center of the crosses mark the posterior median estimates, while the horizontal and vertical lines represent the 95% HDI. Black markers correspond to the CRS or hM4D groups, whereas gray markers denote the control group. Panels (A, B) show results from the CRS experiment and panels (C, D) represent the mPFC inhibition experiment. (E-H) Simulated struggling durations with the PH+rho model. These heatmaps follow the same format as (A–D) and depict total simulated struggling times generated by the PH+rho model. Panels (E, F) show results from the CRS experiment, and panels (G, H) correspond to the mPFC inhibition experiment. (I-L) Explanatory power (r²) of model parameters in predicting cumulative swimming and struggling times. Panels (I, J) depict predictions for cumulative swimming time, while panels (K, L) show predictions for cumulative struggling time. Black curves represent models incorporating all parameters, and colored curves illustrate the contributions of individual parameters. Learning rate-related parameters (α, κ) are most influential during early stages, whereas parameters related to consequence sensitivity (lb, ub, ρ) become increasingly important over time.

To further clarify how these parameters influence behavior over time, we used linear regression models to predict cumulative swimming or struggling durations from parameters sampled from the posterior distributions of the winning models. We then compared each parameter’s explanatory power (r²) at each timestep. This analysis revealed that different parameters exerted varying degrees of influence as the test progresses. In the FST, α from the RW+ada model was the strongest predictor of cumulative swimming time early on, but its influence declined in later stages, whereas lb and ub became more important (Figure 6I, J). In the TST, κ from the PH+rho model was the dominant factor early in the test, while ρ gained significance during later stages (Figure 6K, L). These results were further proofed by a munite-by-munite analysis of each model parameter’s r^2^ (Figure S5).

These findings suggest that conventional analytical approaches, which often focus on late-phase data, may underestimate the impact of learning rate-related factors and overemphasize consequence sensitivity when interpreting rodent behavior.

## Discussion

In this study, we introduced a open-source tool for annotating animal behavior, enabling the efficient collection of detailed behavioral trajectories in the FST and TST. Our computational modeling analysis revealed that both tasks are driven by reinforcement learning principle involving learning, consequence perception, and decision-making. However, the underlying cognitive processes differ between the FST and TST, challenging the common assumption that these tasks are equivalent or mutually validating. By examining model parameters that represent different cognitive components, we found that CRS and mPFC inhibition predominantly increased immobility by altering the learning rates and consequence sensitivity. Additionally, our analysis showed that the impact of these factors varies across different phases of the tests. Consequently, traditional approaches that overlook early-phase data may underestimate the influence of learning-related factors and overemphasizing consequence sensitivity. Overall, our results underscore that detailed behavioral trajectories offer a rich source of information; when examined through computational modeling, they can illuminate the complex cognitive processes driving behavior.

### Facilitating behavioral trajectories analysis with SST

Conventional metrics typically focus on immobility time as a single measure, thus fail to capture the complexity and evolution of behavior. This limitation hinders our understanding of the cognitive processes involved, emphasizing the need for fine-grained behavioral trajectory data. To address these challenges, we developed a Python-based, user-friendly tool that automatically tracks behavioral trajectories in the FST and TST. By providing frame-level temporal resolution, SST enables researchers to identify animal action states with high precision. Such detailed tracking allows the detection of subtle behavioral changes, supports the application of computational models, and fosters deeper insights into the cognitive processes underlying animal behavior.

### Prior FST exposure does not alter behavioral results or winning model in the TST

To directly address the potential confounding effect of prior FST exposure on subsequent TST performance, we conducted a control experiment using naive mice. A total of 45 mice were randomly assigned to two groups: one group underwent only the TST (n = 23), while the other group performed the FST followed by the TST two days later (n = 22). Our analysis showed no significant difference in total immobility time or immobility time during the final 4 minutes between the two groups, and the behavioral trajectories were also comparable (Figure S3), indicating that prior FST exposure did not substantially affect the behavioral response.

To further examine whether prior FST experience might affect underlying cognitive strategies, we applied the same eight reinforcement learning models used in our main analysis to fit the TST behavioral trajectories. Results showed that in both groups, regardless of whether the mice had been exposed to the FST, the same model, PH+rho (dynamic learning rate with consequence sensitivity), consistently emerged as the winning model (Figure S4). This indicates that prior FST exposure did not alter the cognitive processes engaged during the TST. There are two possible explanations for these findings. First, a two-day interval appears sufficient for mice to recover from the transient stress induced by the FST. Second, as discussed before, while the FST and TST yield similar behavioral phenotypes, the cognitive processes underlying each test are not identical. Therefore, performing the FST does not necessarily influence behavior or winning model in the subsequent TST. Taken together, these results support the validity of using both tests in the same animals, provided an adequate recovery period is included.

### Incorporating adaptation of consequence sensitivity in reinforcement learning models

This study marks a pioneering effort to analyze mouse behavior in the FST and TST using reinforcement learning models, providing novel insights into the cognitive processes at work. Traditional models, such as Rescorla-Wagner (RW) and Pearce-Hall (PH), predict a gradual, near-asymptotic decline in the probability of swimming or struggling. However, our FST observations revealed a more dynamic pattern—rather than stabilizing in the later stages, mouse behavior exhibited rebounds and fluctuations, with swimming occasionally reemerging after prolonged immobility. To better capture this complexity, we integrated an adaptation of consequence sensitivity into our models. This adaptation comprises two components: positive adaptation, which alleviates negative subjective experiences after prolonged immobility, and negative adaptation, which amplifies the negative experiences following repeated failures in swimming or struggling. Models incorporating this adaptive factor emerged as the winning models for the FST, accurately predicting immobility durations and capturing the dynamic evolution of behavior. These findings highlight the utility of incorporating adaptive processes into computational models for more accurate simulations of behavioral patterns.

### The FST and TST involve different cognitive processes

Although these tests share broadly similar methodologies, our findings indicate that the models best fitting FST data differ from those best fitting TST data, suggesting that behavior in each task is driven by different cognitive processes. This result calls into question the practice of using the FST and TST interchangeably for cross-validation. The FST appears to emphasize how animals dynamically perceive the consequences of their actions, whereas the TST may be more focused on how attention modulates learning. This distinction could account for the inconsistent results often reported when employing these tests to measure depression-like behaviors. Our findings point to the need for a more nuanced understanding of the cognitive processes that each task probes and underscore the importance of developing more tailored methods for cross-validation.

### Differential effects of CRS and mPFC inhibition on the induction of depressive-like behaviors

Compared to controls, CRS significantly increased immobility in both the FST and TST. In contrast, inhibition of the mPFC resulted in a significant increase in immobility only in the FST, and to a lesser extent than CRS. Computational modeling revealed that both CRS and mPFC inhibition reduced the learning rate (α), amplified behavioral despair following failed escape attempts (reflected in a downward-shifted lower bound of adaptation lb), and accelerated the progression of this despair (indexed by a higher adaptation rate β) during the FST. Notably, only mPFC inhibition increased decision determinism (i.e., sensitivity to value differences, captured by the inverse temperature τ), a change not observed under CRS. In the TST, both manipulations reduced decision determinism (τ); however, only CRS led to a negatively biased evaluation of struggle outcomes (ρ), along with elevated values for the scaling factor (κ) and attention updating rate (η), suggesting heightened sensitivity to aversive outcomes and enhanced learning from negative feedback. Together, these findings suggest that CRS and mPFC inhibition may influence overlapping yet partially distinct cognitive processes that contribute to depressive-like behaviors.

### Limitation of traditional methods: overlooking learning and overemphasizing consequence sensitivity

Our results highlight the critical role of learning in the early stages of the tests, whereas consequence sensitivity takes on greater prominence in later stages. By excluding the first two minutes, a common practice, traditional analyses risks underestimating the importance of learning, while overemphasizing the roles of consequence sensitivity. This oversight may impede our comprehension of depression-like behaviors and limit the development of effective antidepressant treatments. Consequently, we recommend that future research examine the entire behavioral trajectory and integrate computational models to achieve a more comprehensive understanding of the cognitive processes that underlie these behaviors.

### Limitations of the study

While reinforcement learning models offer a valuable framework for interpreting behavioral trajectories, alternative neurobiological mechanisms likely contribute to the observed changes. Acute and chronic stress can induce neuroendocrine and neuromodulatory shifts that influence behavior independently of associative learning. Specifically, acute stress activate the HPA axis and leads to corticosterone release, which promotes passive coping (e.g., increased immobility) via mineralocorticoid (MR) and glucocorticoid receptors (GR)^62–65^. These receptors modulate emotional and motivational processing, suggesting that corticosterone may bias behavior toward immobility without the need for experience-dependent learning. Concurrently, phasic activity of locus coeruleus noradrenergic neurons, triggered by stress, is linked to enhanced struggling in early test phases^66,67^. At the circuit level, stress may induce a shift in control from active to passive coping networks. The medial prefrontal cortex (mPFC) modulates immobility by inhibiting the ventrolateral periaqueductal gray (vlPAG) via the av-BNST^57^. Reduced top-down control may disinhibit vlPAG and increase immobility. In contrast, the VTA-striatal pathway promotes active coping^68^, and behavioral transitions may reflect rebalancing between these competing systems.

Together, these alternative perspectives underscore that the behavioral dynamics observed in the FST and TST likely emerge from an interplay between learning-based processes and state-dependent neuromodulatory and circuit-level mechanisms. While RL models provide a valuable lens for quantifying behavior, a comprehensive interpretation should integrate both computational and biological insights.

## Resource availability

### Lead contact

Requests for information and resources should be directed to the lead contact, Zhihan Li.

### Materials availability

This study did not generate new unique reagents.

### Data and code availability

The source code for SwimStruggleTracker, the behavioral annotation tool developed in this study, is publicly available at: https://github.com/UsamiSumireko/SwimStruggleTracker. Any additional information is available from the lead contact upon request.

## Acknowledgements

This work was supported by the National Natural Science Foundation of China (No. 32300853) and the STI2030-Major Projects (No. 2022ZD0208200).

## Author contributions

Z.L., and Y.L. conceived the project. Z.L., T.L., J.Y., and X.Z. performed the original experiments. Z.L., and T.L developed the tool for automated behavioral recognition and computational models, designed and performed analysis. Z.L., T.L., and Y.L. discussed the results and wrote the manuscript.

## Declaration of generative AI and AI-assisted technologies

The authors utilized LLMs to enhance the readability and language of this work. Following its use, the authors thoroughly reviewed and edited the content as necessary and take full responsibility for the content of the publication.

## Supplemental information

Figures S1–S5, Video S1. Comparison of SST and SMART v3.0 annotations in the TST, related to Figure 2.

## Methods

### Subjects

Male C57Bl/6 mice (weighing 20–30g, 8 weeks old at the start of experiments) were obtained from Beijing SPF Animal Technology Co. for all experiments. The mice were housed under controlled conditions: a 12:12-hour light-dark cycle (lights on at 7:00 AM), a temperature of 25 ± 2°C, and humidity levels of 50 ± 10%. Behavioral experiments commenced after one week of acclimatization and were conducted.

### Chronic restraint stress exposure

The CRS procedure followed a widely accepted paradigm with slight modifications. In brief, mice were restrained in tubes with limited movement for 8 hours daily (10:00 AM–6:00 PM) over 21 consecutive days. Control mice were kept in their home cages without exposure to stress. After each restraint session, the mice were released and returned to their home cages. All experimental procedures were approved by the Institutional Animal Care and Use Committee (IACUC) of the Beijing Institute of Basic Medical Sciences (Approval number: IACUC-06-2019-002).

### Chemogenetic inhibition of medial prefrontal cortex (mPFC) pyramidal neurons

Stereotaxic surgeries were performed under general anesthesia with 1% sodium pentobarbital using a stereotaxic apparatus (RWD Life Science Co., Ltd, Shenzhen, China). To inhibit pyramidal neurons in the mPFC, 500nL of rAAV-CaMKIIα-hM4D(Gi)-mCherry (BrainVTA Co., Ltd, Wuhan, China) was bilaterally injected into the mPFC (250nL for each side) of C57Bl/6 mice at the coordinates: AP +2.34 mm, ML ±0.3 mm, and DV −1.8 mm from Bregma. The control group received the same injection protocol but with rAAV-CaMKIIα-mCherry (BrainVTA Co., Ltd, Wuhan, China). The infusion was performed using a glass pipette connected to a Hamilton syringe fitted with a 33-gauge needle. The syringe was mounted on a syringe pump (RWD Life Science Co.) to ensure precise control over the infusion process. The infusion rate was set to 50 nl/min, allowing for consistent and accurate delivery of the solution. The injection pipette was slowly withdrawn 10 minutes after the infusion. Behavioral experiments were conducted 4 weeks post-surgery. On the day of the testing, Clozapine N-oxide (CNO) was administered intraperitoneally to inhibit pyramidal neurons(5mg/kg), and behavioral assessments began 90 minutes later.

### Behavioral Tasks

The FST was performed in a well-lit room. Mice were individually placed in water-filled cylinders (35 cm in height ×10 cm in diameter) with water maintained at 24-25°C and at depth sufficient to prevent their limbs from touching the bottom. Each mouse was allowed to swim freely for 6 minutes, and its movements were recorded on video.

For the TST, mice were gently removed from their cages, and their tails were secured with adhesive tape approximately 1 cm from the tip. The mice were then suspended by their tails at a height of about 30 cm from the ground, placing them in an inverted position. The test also lasted for 6 minutes, and video recordings ware taken throughout.

### Histology and microscopy

Following the final day of the behavioral experiments, mice were injected with 0.2 mL of a urethane solution at a concentration of 0.3 mg/mL to achieve anesthesia. Subsequently, they underwent transcardial perfusion with 4% paraformaldehyde in PBS for fixation. The brains were then post-fixed overnight, and coronal brain sections (20 μm) were prepared. Cell samples were imaged using a fluorescent microscope, and images were analyzed using ImageJ Pro Plus Version 6.0 (Bethesda, Maryland, USA).

### The Swim Struggle Tracker (SST)

In brief, the SST consists of four modules: the calibration and region of interest (ROI) selection module, the passive motion compensation module, the active motion detection module, and the smooth and thresholding analysis module. We briefly describe the function of each module below.

1. *Calibration and ROI selection module:* As shown in Figure 2A, we mitigate the impact of camera angles on recognition accuracy by applying a perspective transformation to the original images. Prior to detection, the user manually selects the coordinates of the experimental box’s vertices in the video frame. By comparing these coordinates with pre-measured box dimensions, we calculate and apply a perspective transformation matrix to correct the original video frames. On the corrected frames, the largest possible movement area of the mouse is defined as the ROI. The SST offers two options for selecting the ROI: manually adjusting the size by dragging the mouse or choosing from preset sizes. The selected ROI is resized to 360×480 pixels for further analysis.
2. *Passive motion compensation module (optional):* As illustrated in the dashed box in Figure 2B, this module focuses on analyzing the animal’s contour map extracted from the ROI, along with key points required for estimating optical flow. Optical flow techniques are used to track these key points in successive frames. In cases where the animal is passively moving, such as during inertial drift or pendulum motion, its posture remains static. Consequently, the relative positions of key points between adjacent frames remain constant, indicating only an overall shift. The pyramid Lucas-Kanade optical flow algorithm is applied to detect corresponding key points in the following frame. By minimizing the geometric distance between predicted and actual key point positions, the algorithm determines the geometric transformation that best represents the animal’s overall movement. This fitted transformation models the passive motion of the animal. Using this module, the current frame is adjusted to generate a passive motion-compensated prediction, enabling direct comparison with the subsequent frame for further analysis.
3. *Active motion detection module:* As shown in Figure 2B, active movement by the animal results in significant differences between adjacent frames, even after passive motion compensation. We use a three-frame differencing method to calculate inter-frame differences based on the ROI images from frames t and t+1. These differences are then applied a threshold to generate a binary difference image at frame t, which is combined with a cached binary difference image from frame t-1 using intersection operations. Morphological operations (erosion and dilation) are adopted to remove noise, and the processed image is then summed and thresholded to determine the movement state (struggling or immobilize). Figure 2E displays the current video frame, contour detection results (if passive motion compensation is enabled), and combined inter-frame differences to illustrate the tool’s effectiveness.
4. *Behavioral trajectory smoothing and binning module:* As illustrated in Figure 2C, the original behavioral trajectory for each frame often contains significant noise. For example, in a video where a mouse struggles for the first 6 seconds and remains immobile for the following 6 seconds, initial recognition may produce noisy results, as shown in the top row. To address this, Gaussian smoothing is applied to reduce noise and produce a clearer trajectory, as depicted in the second row. Next, thresholding is used to convert the smoothed data into a binary trajectory, shown in the second row from the bottom. Finally, the data is processed through binning and an additional thresholding step, resulting in a behavior trajectory with 2-second bins. This final trajectory, shown in the bottom row, is both more accurate and robust for subsequent analysis.

We validated SST’s performance by comparing its results to human expert annotations. Significant correlation was observed in immobility times (Figure 2D, FST: r = 0.411, p = 0.002; TST: r = 0.823, p < 10^−5^, Pearson correlation), demonstrating that the tool’s outputs closely align with manual annotations.

### Reinforcement learning models

We implemented 8 reinforcement learning models to explore different value updating mechanisms and cognitive processes underlying behaviors in FST and TST.

1. *Rescorla-Wagner (RW) model*: The Rescorla-Wagner model is a classic model of error-driven predictive learning. It assumes that mice update their beliefs by incorporating a fixed proportion of prediction error. Given that swimming and struggling deplete the mice’s physical energy, making the subsequent failing to escape experiences more unpleasant than remaining immobile, we encoded the consequence of swimming and struggling as −1, while immobility was encoded as 0. The predictive error at timestep t was calculated as follows:

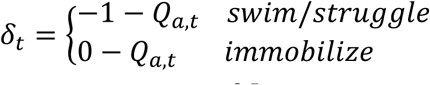 The value expectation was updated as:

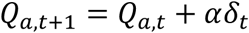 Here, 𝛿_𝑡_represents the prediction error at timestep t, while 𝑄_𝑎,𝑡_and 𝑄_𝑎,𝑡−1_represent the value expectations for the actions taken by the mice at timesteps t and t-1, respectively. The parameter α is the learning rate, which governs the extent to which the prediction error is integrated into the updated value expectation. A sigmoid function was applied to compute the probability of swimming/struggling based on the values expectation of actions. At each trial t, the probability of swimming/struggling 𝑝_𝑠,𝑡_was given by:

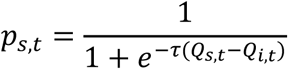 In this equation, 𝑄_𝑠,𝑡_ and 𝑄_𝑖,𝑡_ represent the value expectations of swimming/struggling and immobility, respectively. The probability of immobility is 1−𝑝_𝑠,𝑡_. This decision function was adopted by RW, RW+rho, RW+ada, PH, PH+rho, and PH+ada models.
2. *RW model with perseveration (RW+per)*: Building on the RW model, this model incorporates the influence of behavioral perseveration on the decision-making process. Behavioral perseveration is modeled based on the frequency of unchosen actions 𝑢𝑐𝑡_𝑎,𝑡_, which is updated according to the following rule:

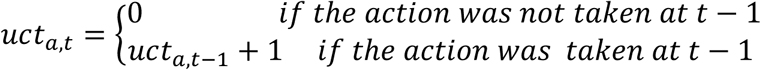 The 𝑢𝑐𝑡_𝑎,𝑡_are considered in decision. At each trial t, the probability of swimming/struggling 𝑝_𝑠,𝑡_ was given by:

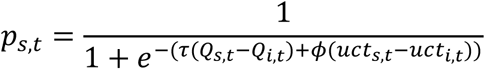

Where 𝑢𝑐𝑡_𝑠,𝑡_and 𝑢𝑐𝑡_𝑖,𝑡_refers to the unchosen time for swimming/struggling and immobile, respectively. 𝜙 refers to the importance of perseveration.
3. *RW model with consequence sensitivity (RW+rho)*: Building on the RW learning rule, this model incorporates the sensitivity to the consequence of swimming and struggling. The sensitivity to the consequence is represented by the parameter ρ. In this model, after swimming or struggling, the predictive error is computed as follows:

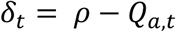 This adjustment reflects the idea that the mice perceive the consequences of their actions (e.g., swimming and struggling) with varying degrees of sensitivity, as captured by 𝜌.
4. *RW model with consequence sensitivity adaptation (RW+ada)*: This model further extends the RW learning rule by accounting for the adaptation of the mice’s sensitivity to the consequence of actions. We hypothesize that repeated failed escape attempts through swimming or struggling or after prolonged reliance on immobility lead to changes in perceiving consequence of actions. The adaptation process is described as:

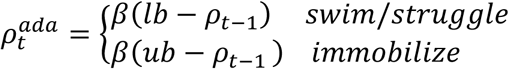 Here, the adapted consequence sensitivity 𝜌_t_^𝑎𝑑𝑎^replaces the fixed parameter ρ from the previous model, with 𝛽 representing the rate of adaptation. The lower and upper bounds of sensitivity adaptation are represented by parameters 𝑙𝑏 and 𝑢𝑏, respectively. The prediction error is then updated as:

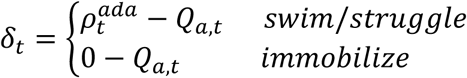
5. *Pearce-Hall (PH) Model*: Unlike the Rescorla-Wagner rule, which assumes a constant learning rate, the Pearce-Hall model dynamically adjusts the learning rate based on the level of surprise or unpredictability. The value expectation is updated as follows:

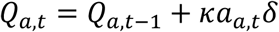 Here, 𝑎_𝑎,𝑡_represents the attention allocated to the action at time t, 𝜅 is a normalization parameter for learning rate. The attention 𝛼_𝑎,𝑡_is updated based on the prediction error 𝛿_𝑡_as follows:

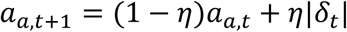

Where 𝜂 refers to the updating rate for attention.
6. *PH model with perseveration (PH+per)*: Combines the PH learning rule with perseveration, as defined in the RW+per model.
7. *PH model with consequence sensitivity (PH+rho)*: Combines the PH learning rule with consequence sensitivity, as defined in the RW+rho model.
8. *PH Model with consequence sensitivity adaptation (PH+ada)*: Combines the PH learning rule with consequence sensitivity adaptation, as defined in the RW+ada model.

### Hierarchical Bayesian models

We used a hierarchical Bayesian framework to model individual behavioral trajectories. This approach allowed us to estimate participant-specific parameters while simultaneously leveraging data from the entire experimental group to improve estimation precision. Specifically, individual parameters were drawn from group-level distributions as follows:

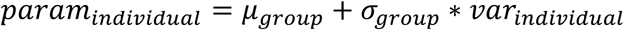

Here, 𝑝𝑎𝑟𝑎𝑚_𝑖𝑛𝑑𝑖𝑣𝑖𝑑𝑢𝑎𝑙_ represents the parameter for each individual mouse, 𝜇_𝑔𝑟𝑜𝑢𝑝_ and 𝜎_𝑔𝑟𝑜𝑢𝑝_ are the group-level mean and standard deviation of the parameter, respectively, and 𝑣𝑎𝑟_𝑖𝑛𝑑𝑖𝑣𝑖𝑑𝑢𝑎𝑙_ refers to the individual-level variance. This hierarchical structure effectively captures both individual differences and shared underlying processes, offering a robust and nuanced representation of the observed behavioral dynamics.

### Parameter estimation and model selection/validation

Parameter estimation was conducted using hierarchical Bayesian analysis (HBA) implemented in the Stan language^69^ via Python (cmdstanpy^70^). Posterior inference was performed through Markov chain Monte Carlo (MCMC) sampling. The models were fitted separately for each and each test. The Gelman-Rubin convergence diagnostic (R-hat) was applied to assess MCMC stability. With all R-hat statistics falling within the recommended threshold of 1.05, the chains exhibited sufficient mixing and stationarity, supporting the reliability of posterior estimates.

Model comparison was used to evaluate how well the models fit the data, accounting for model complexity. Model comparison was assessed using Bayesian bootstrap and model averaging, wherein the log-likelihoods of each model were evaluated based on the posterior simulations, and a weight was assigned to each model. The model weights included a penalization term for complexity and a normalization term relative to the number of models being compared. For each group, the model weights summed to 1. The model with the highest weight was defined as the “winning model,” indicating that it best fit the observed behavioral data of the mice.

To perform a posterior predictive check, we simulated behavioral trajectories for each animal in each experimental manipulations and test, based on the parameters of the winning model. Specifically, the model generated random draws from each animal’s joint posterior distribution to predict actions. This process was repeated 2000 times per animal for each test. We compared each animal’s synthetic data to their experimental observations, focusing on the immobility times both throughout the test and in the final 4 minutes, the first immobile time, and the behavioral trajectories of mice.

### Analysis of trends in research publications

To examine the research trends involving the FST and TST, the annual number of publications from 1995 to 2024 was collected from the Web of Science database. The analysis followed a two-step query process to ensure specificity to studies related to affective and stress-related disorders as well as the FST and TST.

The initial query used the following terms to identify publications related to affective and stress-related disorders:

TS = ((depression) OR (depressed) OR (depressive) OR (dysthymia) OR (“affective disorder”) OR (“mood disorder”) OR (anhedonia) OR (despair) OR (helplessness) OR (anxiety) OR (stress) OR (“social defeat”)).

This search was refined by adding terms specific to the FST or TST.

For the FST, the query was:

TS = ((FST) OR (“Forced Swimming Test”) OR (“Forced Swim Test”)). For the TST, the query was:

TS = ((TST) OR (“tail suspension test”) OR (“suspension test”) OR (tail-hanging test)).

The results from each refined search were categorized by year of publication to analyze annual trends. The data were visualized as a bar graph to illustrate the temporal progression of research activity in the field.

### Statistical Analysis

Comparative statistical tests, correlation analyses, and linear regressions were conducted in Python using libraries such as SciPy^71^ and scikit-learn^72^. Mann-Whitney U tests were employed to compare immobility times and model parameters between groups. Pearson correlation analyses were used to examine relationships between each individual’s posterior predictive mean of immobility time, latency to immobility, the corresponding experimental observations, and the agreement of immobility times identified by SST compared to human annotations.

## Footnotes

1 https://github.com/UsamiSumireko/SwimStruggleTracker

