## Supplemental Figures for "Beyond Immobility: Computational Modeling Reveals Cognitive Processes in Simple Rodent Depression Tests"

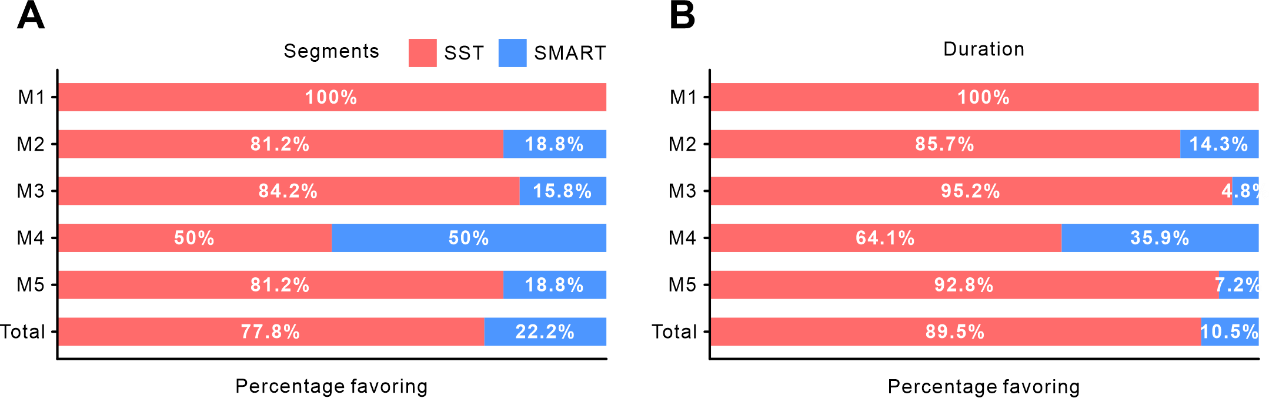

**Figure S1. Human expert evaluation of annotation discrepancies between SST and SMART v3.0 on a TST video. The TST video containing five mice was analyzed using both SST and SMART v3.0, with behavioral annotations scored in non-overlapping 2-second windows to classify each animal’s dominant behavior as either struggling or immobile. In cases where the two systems produced conflicting annotations, human experts adjudicated which annotation was more accurate. (A) Proportion of segments (i.e., contiguous time windows) in which experts favored SST (red) or SMART (blue). (B) Expert preferences based on the total duration of all disputed time windows. M1–M5 denote the five individual mice.**

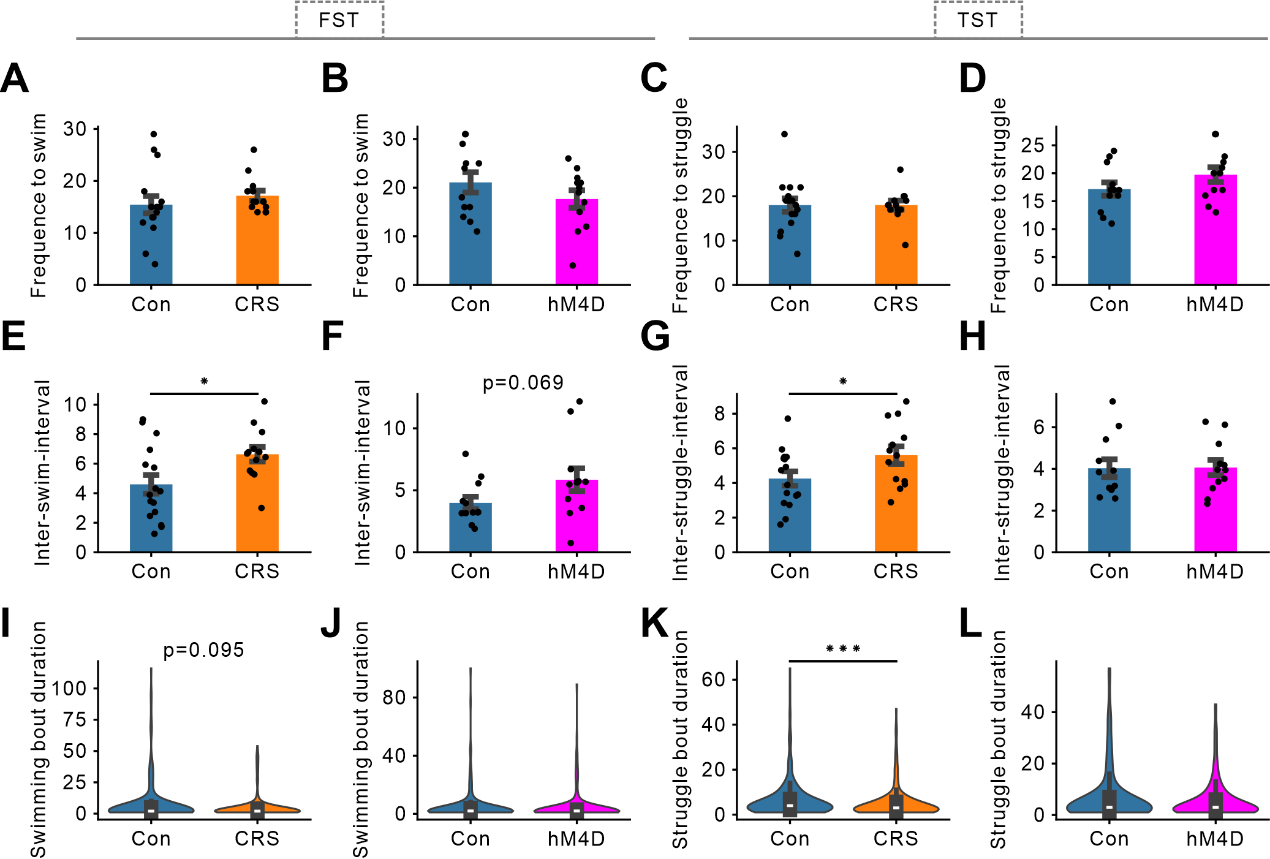

**Figure S2. Frequency to struggle, inter-struggle-interval, and struggle bout duration in the FST and TST. (A–D) Frequency to swim/struggle in the FST (A, B) and TST (C, D). Blue, orange, and magenta bars represent the control, CRS, and hM4Di groups, respectively. Each black dot denotes an individual animal. Error bars indicate the standard error of the mean (SEM). (E–H) Inter-struggle interval, defined as the time between consecutive swim/struggle bouts, in the FST and TST. Plot format is consistent with panels (A–D). (I–L) Struggle bout duration, defined as the length of each continuous episode of swimming or struggling, in the FST and TST. Data are presented in the same format as in panels (A–D). Asterisks denote significant differences from the controls as determined by the Mann–Whitney U test; ***p < 0.001, *p < 0.05.**

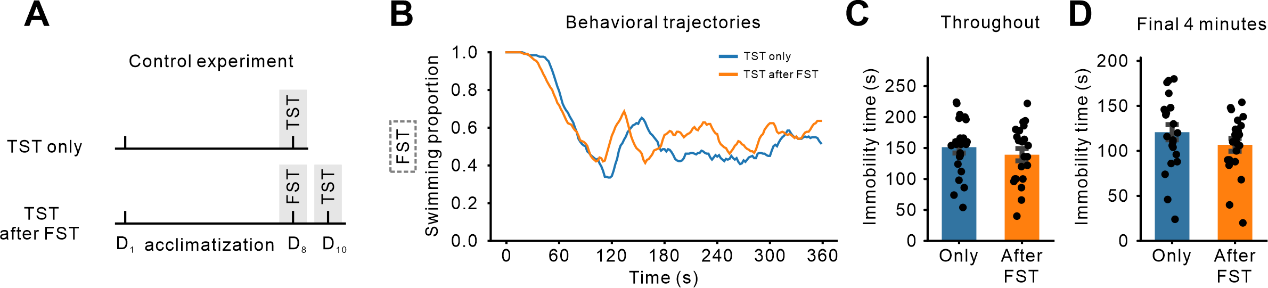

**Figure S3. Prior FST exposure does not significantly alter subsequent TST behavior. (A) Experimental design: 45 mice were randomly assigned to two groups. A TST-only group (n=23) and a group that performed the FST followed by the TST two days later (TST after FST, n=22). (B) Averaged behavioral trajectories during the TST. The graph shows no substantial differences in the evolution of behavior over time between the two groups. (C, D) Total immobility time for the entire TST session (C) and the final 4 minutes (D). No significant differences were observed between the two groups (Mann-Whitney U test, total immobility time: p=0.440; final 4 minutes: p=0.216). Blue and orange bars represent the TST-only and TST after FST groups, respectively. Black dots indicate individual animals. Error bars represent the standard error of the mean (SEM).**

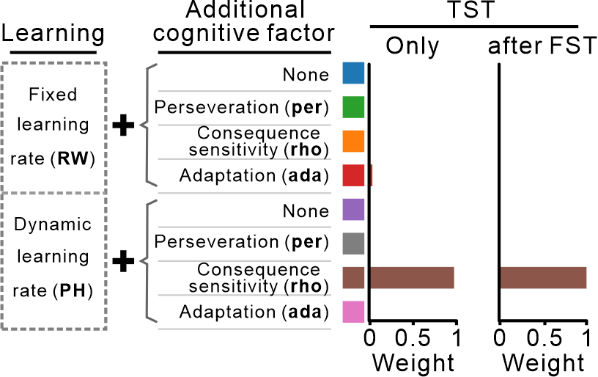

**Figure S4. Prior exposure to the FST does not alter the winning model in the TST. We applied our candidate reinforcement learning models to fit the behavioral trajectories from the TST-only group and the TST after FST group. Model comparison was conducted by computing model weights, which represent the relative likelihood of each model being the true generative process. The model with the highest weight was identified as the winning model. As shown by the consistent brown bars, the PH+rho model (dynamic learning rate with consequence sensitivity) emerged as the winning model in both conditions, indicating that prior FST exposure did not alter the cognitive processes engaged during the TST.**

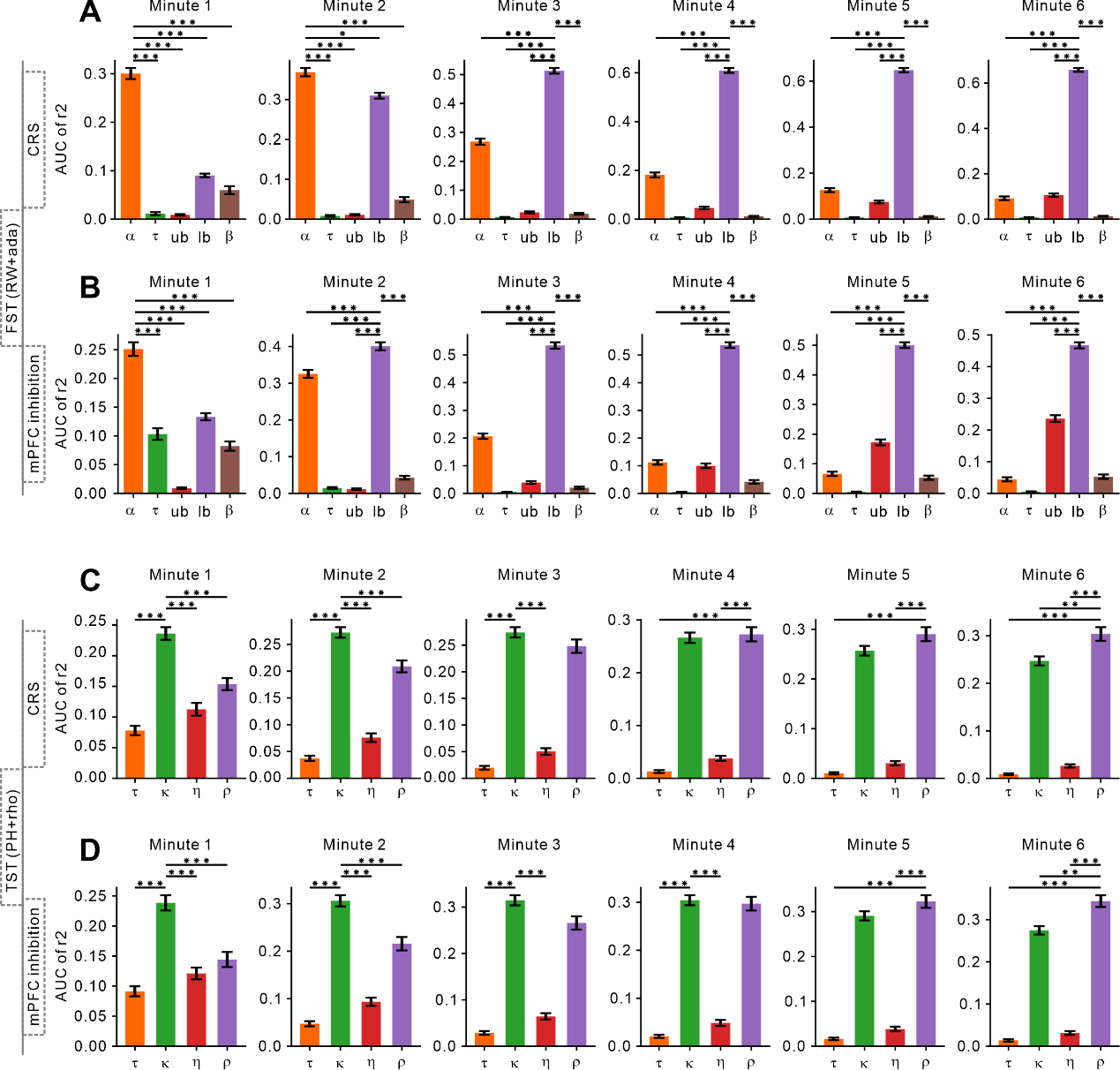

Figure S5. Minute-by-minute explanatory power (r²) of model parameters for behavioral prediction. (A, B) Explanatory power of individual parameters from the RW+ada model in predicting cumulative swimming time during the FST under CRS (A) and mPFC inhibition (B) conditions. Behavior was analyzed in six consecutive 1-minute bins. Colors indicate specific parameters: orange = learning rate (α), green = inverse temperature (τ), red = upper bound of adaptation (ub), purple = lower bound of adaptation (lb), brown = adaptation rate (β), Error bars indicate SEM. (C, D) Explanatory power of parameters from the PH+rho model in predicting cumulative struggling time during the TST under CRS (C) and mPFC inhibition (D), using the same minute-by-minute analysis. Bar colors: orange = inverse temperature (τ), green = scaling factor (κ), red = attention updating rate (η), purple = consequence sensitivity (ρ). Due to violations of normality, Friedman tests were applied to assess overall differences in explanatory power across parameters within each time bin. Post hoc comparisons were conducted using Wilcoxon signed-rank tests with Bonferroni correction. Asterisks indicate statistically significant differences between the most predictive parameter and others in each minute (***p < 0.001, **p < 0.01, *p < 0.05).
